# Association between Prevalent Flagellated Bacteria in Gut Microbiota and Cardiovascular Benefits in Humans

**DOI:** 10.1101/2023.05.04.539500

**Authors:** Jensen H. C. Yiu, Jieling Cai, Samson W. M. Cheung, Karie Tsz-Ching Chin, Chi Fai Chan, Edward S.C. Ma, Rakesh Sharma, Bernhard Dorweiler, Connie W. Woo

## Abstract

**Background:** Increasing evidence has shown associations between gut microbiota and cardiovascular risks. However, whether sex influences the cardiovascular outcome of gut microbiota remains elusive.

**Methods:** The gut microbiota metagenomic data from two previous population-based cohorts and the proteomics data from human liver samples were analyzed.

**Results:** Through a sex-based analysis of 500 Functional Genomics Project (500FG) cohort, we found that the capacity of producing flagellin in *Eubacterium rectale, Roseburia Intestinalis* and *Roseburia inulinivorans* partially explained the levels of high-density lipoprotein-cholesterol (HDL-C) and apolipoprotein A1 in female participants. By comparing the bacterial species showing positive correlations with HDL-C and the flagellin species found in human liver samples, we found that *E. rectale* and *R. inulinivorans* appeared to be the major prevalent flagellated species in gut microbiota contributed to the host’s HDL-C level. The analysis of the Chinese-based cohort in which the male participants had relatively higher abundance of these two bacteria, the abovementioned positive correlation was also observed.

**Conclusions:** The flagellin-producing function of *E. rectale* and *R. inulinivorans* in the gut microbiota partially explained the HDL-C level in the host, and there is a sex-specific difference in the sensitivity of this beneficial interaction. Our findings suggest a potential interaction between commensal bacteria and the host that yields a cardiovascular benefit.

**Clinical Perspective:** *What is new?:*

- The flagellin-producing capacity of certain species in gut microbiota is positively associated with the HDL-C and ApoA1 levels in human.
- By comparing the flagellated bacteria in the gut and flagellin species in the liver, *Eubacterium rectale* and *Roseburia inulinivorans* are shown to be the most prevalent species contributing to such cardiovascular benefits.
- The sensitivity of such beneficial interaction with the gut flagellated bacteria is different between male and female.

*What are the clinical implications? (maximum 100 words, formatted as 2-3 bullets):*

- Beside the role in metabolic inflammation, gut microbiota can be beneficial against cardiovascular risk by modulating HDL-C level through certain flagellated species.
- The interaction between flagellated bacteria in the gut and the host provide a different insight in exploring potential therapeutic targets to increase HDL-C level.

## Introduction

Men and women have distinctive lipid profiles and women are known to be protected against cardiovascular disease because of their higher HDL-C level. Women, especially at the premenopausal state, are often excluded or underrepresented in cardiovascular studies. The decrease in estrogen thus far is the only blame for postmenopausal women losing such sex-specific benefit. Since increasing evidence has shown associations between gut microbiota and cardiovascular risks^1–6^ and distinctive patterns of gut microbiota between males and females^7,8^, we wonder whether gut microbiota also contributes to the sex difference in the cardiovascular benefits. A population-based cohort studies suggested that gut microbiome were correlated with body mass index (BMI), triglyceride and HDL-C levels.^9^ HDL-C level was found positively correlated with species richness in a study focusing on the connections between habitual diet, gut microbiomes and cardiometabolic markers.^10^ However, these studies have yet to address any sex difference or what kind of properties of the gut microbiota contributing to such correlations.

Our previous murine study showed that high-fat diet increased the abundance of flagellated bacteria in the gut, and the flagellin released by these bacteria activated hepatic toll-like receptor 5 (TLR5), resulting in elevation of ApoA1 and HDL-C levels.^11^ Here, we separately analyzed the gut microbiota of females and males from the 500 Functional Genomics (500FG) Project, a population-based cohort, and discovered a positive correlation between the flagella-producing capacity in gut microbiota and HDL-C level in females only. Flagellins derived from gut flagellated bacteria were detected in human liver. *Eubacterium rectale* and *Roseburia inulinivorans* were the two prevalent species flagellins of which were detectable in liver and showing positive correlation between their flagellum-producing capacity and host’s HDL-C level. Moreover, the presence of TLR5 but not NLRC4/NAIP, an inflammasome complex that is also activated by flagellins, in human liver limits the inflammation stimulated by the filtrated flagellins. Although such benefit of gut flagellated bacteria was observed in female only in the 500FG cohort, in a Chinese cohort in which males had relatively higher abundance of *E. rectale* and *R. inulinivorans*, the positive correlation between flagellum-producing capacity and HDL-C level was also observed. Take together, our data support the feasibility of targeting hepatic TLR5 to increase HDL-C level.

## Methods

Please see the Major Resources Table in the Supplemental Materials. The data that support the findings of this study are available from the corresponding authors upon reasonable request.

### Population cohorts

The 500 Functional Genomics (500FG) project was initiated in 2013 and recruited 534 adult healthy volunteers from the Netherlands.^10^ Inclusion criteria were (1) above 18 years of age and (2) Western European descent, and exclusion criteria were (1) pregnancy/breastfeeding, (2) chronic or acute disease at the time of assessment, and (3) use of chronic or acute medication during the last month before the study. Only the samples with shotgun metagenomic data on gut microbiome, complete physiological parameters and lipid profile were included in our study, yielding 347 samples (*n* = 193 females, *n* = 154 males) in the final analyses. The Chinese cohort consisted of 942 healthy participants across 6 ethnicities in urban and rural China.^12^ After excluding participants (1) without HDL-C level measurement and shotgun metagenomic data on gut microbiome, (2) who took antibiotics, Western medicine or traditional Chinese medicine and living in the rural areas, 316 samples (*n* = 213 females, *n* = 103 males) were analyzed in our study.

### Human tissues

Human liver tissues were purchased from BioIVT (France) and free of pathology and infection by HBV/HCV/HIV. Deidentified specimens of retroperitoneal adipose tissues and aortic walls were obtained from patients undergoing open aortic replacement at the Division of Vascular Surgery, University Medical Center, Johannes Gutenberg University (Mainz, Germany). All patients gave informed consent and institutional review board approval was waived (use of excess/discarded material). Specimen were snap-frozen in liquid nitrogen immediately after collection.

### Gut microbiome profiling

#### Quality control and preprocessing

High-quality metagenomic data from the 500FG and the Chinese cohorts were downloaded from Sequence Read Archive of NCBI. The sequence files were first preprocessed using KneadData (Harvard University, USA) with the default settings, including removal of human contaminant sequences (hg37dec_v0.1) identified by bowtie2 using the very-sensitive mode, reads with low-quality reads (Q < 20) and fragmented short reads (<50 bp).

#### Microbial taxonomic and functional profiles

The taxonomic profile was determined using MetaPhlAn 3.0 (Harvard University, USA). This version was built using 99,237 reference genomes representing 16,797 species retrieved from Genbank as of January 2019. The functional profiling was performed using HUMAnN 3.0 (Harvard University, USA). In brief, the cleaned reads were first mapped to clade-specific marker genes to identify community species by MetaPhlAn 3.0, followed by mapping to pangenomes of identified species (ChocoPhlAn database). The unmapped reads were aligned to a non-redundant database, Uniref90 (Version 201901b). All the aligned results were then estimated with total gene family abundance per species and community based on alignment quality, gene length and gene coverage. The functional enrichment analysis was completed by grouping the gene families using Gene Ontology (GO) annotations. Abundances in counts per million (cpm) were analyzed in this study.

### Proteomic identification using LC-MS/MS

Human liver and aortic lesion protein lysates were separated on SDS-PAGE gels and visualized with Coomassie stain. The positions at which flagellins were detected using anti-flagellin antibodies from Covalab Inc. (France) and Abcam Inc. (UK) were excised for flagellin identification, whereas the proteins above 72 kDa were excised for identification of TLRs and NLRs. The gel slices were then subject to in-gel digestion by trypsin followed by LC-MS/MS analysis at Centre for PanorOmic Sciences, HKU. In brief, gel slices were subjected to reduction and alkylation by 10 mM tris(2-carboxyethyl)phosphine (TCEP) and 55 mM 2-chloroacetamide (CAA), respectively following by incubation with trypsin (1 ng/µl) overnight at 37 °C. Subsequent tryptic peptides were extracted from the gel with 50% acetonitrile (ACN)/5% formic acid (FA) and 100% ACN sequentially and desalted. The eluted peptide mixture was loaded onto an Aurora C18 UHPLC column (75 μm i.d. × 25 cm length × 1.6 μm particle size) (IonOpticks, Australia) and separated using a linear gradient of 2-30% of buffer B (0.1% FA in ACN) at a flow rate of 300 nl/min buffer A (0.1% FA and 2% ACN in H_2_O) for 100 min on nanoElute Nano-Flow UHPLC System coupled to timsTOF Pro mass spectrometer (Bruker, USA). MS data was collected over a *m*/*z* range of 100 to 1700, and MS/MS range of 100 to 1700. During MS/MS data collection, each TIMS cycle was 1.1 s and included 1 MS + an average of 10 PASEF MS/MS scans.

#### Sequence identification and validation

Raw mass spectrometry data were processed using MaxQuant 2.0.1.0 (Max Planck Institute for Biochemistry, Germany). For the evaluation of flagellin, raw data were searched against human UniProt FASTA database (Apr 2020) containing 74,824 entries or a customized database containing flagellin proteins identified in the gut microbiota analysis of the 500FG cohort with duplicates removed, using settings as below: oxidized methionine (M), acetylation (Protein N-term) were selected as dynamic modifications, and carbamidomethyl (C) as fixed modifications with minimum peptide length of 7 amino acids was enabled. For the evaluation of TLRs and NLRs, raw data were searched against databases containing all members from TLRs or NLRs, respectively. Confident proteins were identified using a target-decoy approach with a reversed database, strict false discovery rate of 1% at peptide and peptide spectrum matches (PSMs) level with minimum ≥1 unique peptide.^13^ To validate the detected peptides were truly bacterial flagellins, we spiked one of the samples with known amounts of purified flagellin from *Bacillus subtilis*, a species that was not identified in the 500FG cohort, (InvivoGen, USA) followed by proteomic analysis, and the flagellin from *B. subtilis* was detected in the spiked samples. Moreover, the identified flagellin peptides were also subjected to Protein BLAST alignment (National Center for Biotechnology Information, USA) against human proteome for validation (See Supplemental Methods in Supplemental Material).

### Western immunoblotting

Immunoblotting was conducted as described previously.^11^ The anti-flagellin antibodies were purchased from Covalab Inc. and Abcam Inc. The membranes were reblotted with anti-β-actin (Sigma-Aldrich, USA) for detecting differences in protein loading.

### Real-time PCR

Total RNA from tissues was isolated RNeasy Mini Kit (Qiagen). RNA was reverse-transcribed into cDNA using ImProm-II™ Reverse Transcription System (Promega, WI, USA). Real-time PCR was conducted using the SYBR green PCR reagent (Roche) and StepOnePlus™ System (Applied Biosystems, CA, USA). Amplicons were verified using agarose gel electrophoresis and Sanger sequencing. The sequences of the primers are listed in Table S1.

### Statistical analyses

Statistical analyses were performed using R software Version 4.0.5. The difference in gut microbiota pattern was evaluated by the permutational ANOVA (PERMANOVA) using the adonis function from the vegan package in R. Normality was checked using the Shapiro-Wilk test. Spearman partial correlation analysis was performed using the ppcor packing in R and cutoff at the 90^th^ percentile was applied to exclude extreme values. *P* values were adjusted for multiple testing based on the Benjamini-Hochberg method and reported as *q* values. Significant associations were defined by *P* < 0.05 and *q* < 0.2 when multiple testing was applied.

### Data availability

The gut metagenome data are available at NCBI SRA BioProject under accession ID PRJNA319574 (the 500FG cohort) and PRJNA588513 (the Chinese cohort). The lipid and cytokine profiles of the 500FG cohort are available at https://hfgp.bbmri.nl/ while the data for the Chinese cohort are available in the original paper.^12^ Proteomics data have been deposited to the Proteomics Identification Database under the accession ID PXD041941.

### Code availability

Source codes for KneadData, MetaPhlAn 3.0 and HUMAnN 3.0 are available at https://github.com/biobakery/.

## Results

### Sex differences in the correlations between gut microbiota and lipoproteins

We included individuals from the 500FG cohort with shotgun metagenomic sequences on gut microbiome and comprehensive lipid profiles.^14^ After controlling the data quality and completeness, 347 samples (193 females and 154 males) were analyzable. The distributions of age, BMI and the levels of plasma lipids and lipoproteins, and their correlation analyses were listed in Table 1. When we segregated the sexes, significantly higher HDL-C, HDL-TG and LDL-TG levels were observed in females while higher BMI and VLDL-TG level in males (Table 1). Gut microbiota analysis also revealed distinct microbial pattern (PERMANOVA, *P* = 0.001) and intra-sample diversity (α-diversity) in females and males (Figure 1A). We further analyzed the correlations between α-diversity and various plasma lipid parameters in both sexes separately with the adjustment of BMI and age, and in addition, oral contraceptive intake for females because oral contraceptives are known to affect lipoproteins.^15^ In females, the α-diversity was positively correlated with most of the plasma lipoproteins, among which HDL-C level showed the strongest association. By contrast, negative correlations with VLDL-TG and TG levels were observed in males (Figure 1B, Data Set 1). These findings were congruent with a previous study focusing on the connections between habitual diet, gut microbiomes and cardiometabolic markers, in which gut microbiota was suggested to explain the variances in HDL-C and TG levels.^10^ Among the 67 bacteria present in ≥50% of the 500FG cohort, the abundances of 41 strains showed positive correlations with the HDL-C level in females, and *Roseburia intestinalis* and *R. inulinivorans* were the strains with the strongest correlation. Conversely, 49 out of these 67 strains had negative correlations with the TG level in males and *Ruminococcus torques, Dorea formicigenerans* and *Intestinibacter bartlettii* showed the strongest (Figure 1C, Data Set 2). As expected, since TG in circulation is mainly carried by VLDL particles, the pattern of the correlations with VLDL-TG level was similar to that with TG level in males (Figure 1C).

**Table 1.** Comparison of females and males from the 500FG cohort.

| Parameter | Female<br>*n* = 193 |  | Male<br>*n* = 154 |  | *P* value |
| --- | --- | --- | --- | --- | --- |
|  | Mean ± S.D. | Range<br>(90<sup>th</sup> percentile) | Mean ± S.D. | Range<br>(90<sup>th</sup> percentile) |  |
| Age | 26.642 ± 12.005 | 18 – 70<br>(49.6) | 28.33 ± 13.08 | 18 – 70<br>(52.2) | 0.212 |
| BMI (kg/m<sup>2</sup>) | 22.087 ± 2.541 | 17.374 – 33.024<br>(24.964) | 23.434 ± 2.806 | 15.095 – 34.416<br>(26.585) | 4.04 × 10<sup>-6</sup> * |
| HDL-C (mM) | 1.545 ± 1.195 | 0.207 – 9.752<br>(2.437) | 0.901 ± 0.587 | 0.167 – 3.036<br>(1.710) | 2.37 × 10<sup>-10</sup> * |
| LDL-C (mM) | 1.304 ± 1.029 | 0.090 – 8.881<br>(1.989) | 1.174 ± 0.825 | 0.052 – 4.887<br>(2.213) | 0.205 |
| VLDL-C (mM) | 1.201 ± 1.176 | 0.158 – 11.611<br>(2.227) | 1.342 ± 1.012 | 0.127 – 8.156<br>(2.382) | 0.239 |
| TC (mM) | 1.425 ± 1.544 | 0.211 – 16.052<br>(2.430) | 1.128 ± 0.893 | 0.188 – 6.065<br>(2.191) | 0.035 * |
| HDL-TG (mM) | 1.470 ± 1.323 | 0.213 – 8.858<br>(2.723) | 0.980 ± 0.623 | 0.092 – 3.694<br>(1.685) | 0.025 * |
| LDL-TG (mM) | 1.558 ± 1.240 | 0.152 – 10.417<br>(2.900) | 0.907 ± 0.649 | 0.142 – 4.033<br>(1.735) | 1.08 × 10<sup>-9</sup> * |
| VLDL-TG (mM) | 1.119 ± 0.882 | 0.125 – 6.530<br>(2.183) | 1.382 ± 0.976 | 0.222 – 6.294<br>(2.611) | 0.009 * |
| TG (mM) | 1.216 ± 1.045 | 0.181 – 7.252<br>(2.203) | 1.284 ± 0.969 | 0.179 – 6.751<br>(2.528) | 0.537 |

\* *P* < 0.05, unpaired two-tailed *t* test

**Figure 1.**
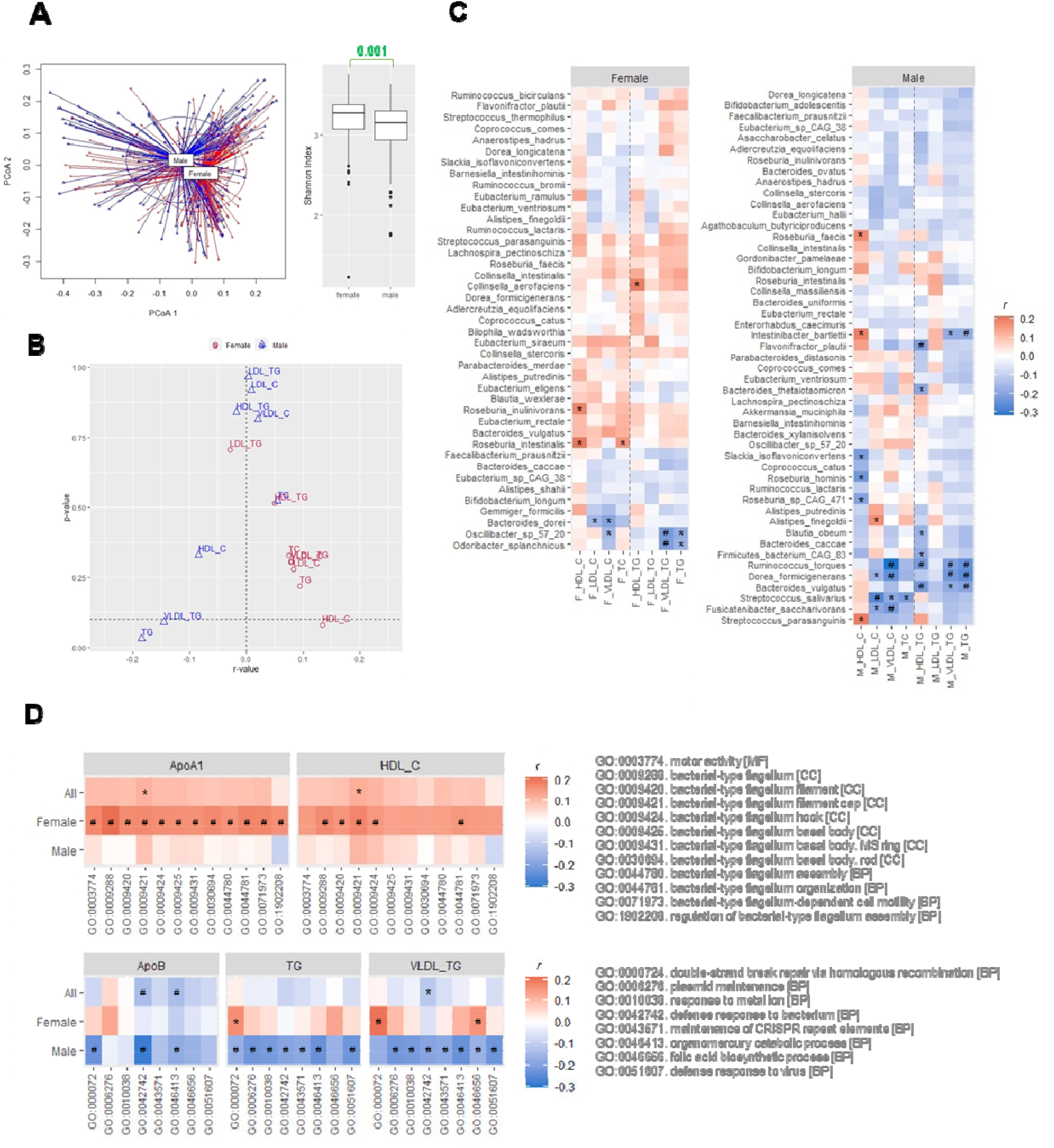
Positive association of microbial diversity with ApoA1 and HDL-C levels in females. (A) β and α diversities evaluated by principal coordinate analysis (PCoA) and Shannon index, respectively, of gut microbiota from female and male subjects of the 500FG cohort. Two-sided unpaired *t* test. (B) Correlations of α diversity with the lipid contents of various types of lipoproteins. (C) Correlations of the abundance of bacteria with the lipid contents of various types of lipoproteins. Species with positive correlations with HDL-C level were selected for the panel of females and those with negative correlations with TG level were selected for the panel of males. (D) Correlations of the flagellum-related pathways with ApoA1 and HDL-C levels and correlations of the survival-related pathways with ApoB, TG and VLDL-TG levels. Spearman correlation adjusted with age and BMI for all, and the use of oral contraceptives for females followed by Benjamini-Hochberg procedure for multiple comparison in (B) to (D). ^*^*P*<0.05, ^#^*P*<0.05 and *q*<0.2. BP, biological process; CC, cellular component; MF, molecular function.

### Association between flagellum-related pathways and HDL-C level in the female of 500FG cohort

Next, we examined the correlations between lipoproteins and the pathways from Gene Ontology (GO) functional enrichment analysis of the gut microbiota. First, we ranked the pathways positively correlated with the HDL-C level by the magnitude of correlation coefficients for females. Among the top 100 pathways, 36 were categorized as biological processes, 11 as cellular components and 53 as molecular functions. Four out of the 36 biological-process pathways and 7 of the 11 cellular-component pathways represented flagellum-related (FLA) pathways (Figure S1, Data Set 3). These aligned with the taxonomical analysis as *Roseburia* spp. are flagellated. We then performed the same for males by ranking pathways negatively associated with TG level instead. In contrast, the top 100 pathways were mainly under the molecular-function ontology and mostly related to bacterial growth and adaptation to habitats (Figure S1, Data Set 3). We further evaluated the correlations of FLA pathways with plasma ApoA1 level, the functional constituent of HDL particles, and the correlations were even stronger than those with HDL-C level in females, particularly the GO:0009288 pathway termed bacterial-type flagellum, which is the topmost cellular-component GO term for bacterial flagella (Figure 1D, Data Set 4). One of the concerns of gut microbiota in the perspective of cardiovascular risks is the potential induction of systemic inflammation, especially when the motility-conferring flagella are often viewed as a virulence factor of bacterial invasion.^16^ Hence, we also evaluated the correlations of FLA pathways and various cytokines. Intriguingly, these pathways consistently showed positive correlations with the level of adiponectin, an adipokine known for its anti-inflammatory effect, in both males and females (Figure S2, Data Set 5). The level of interleukin-18 binding protein (IL18BP), a decoy receptor of the proinflammatory IL18, also showed weak positive correlations in females but negative correlations in males (Figure S2, Data Set 5). By contrast, the proinflammatory cytokines only had weak or no correlations with FLA pathways regardless of sexes (Figure S2, Data Set 5). These data suggest a commensal cardiometabolic benefit between gut flagellated bacteria and the host.

### Detection of flagellins derived from fecal flagellated bacterial species in human liver

Flagellins, the building blocks of bacterial flagella, are the known proteins that interact with the host. As flagellins vary among different strains of bacteria, we wanted to find out which strains of bacteria contributed to the positive correlation between FLA pathways, ApoA1 and HDL-C levels. The FLA pathways derived from *Roseburia intestinalis* and *R. inulinivorans* showed the strongest correlations with ApoA1 and HDL-C levels in female followed by *Eubacterium rectale* and *E. siraeum*, whereas those from *Flavonifractor plautii* and *R. faecis* were more strongly correlated in males (Figure 2A, Data Set 6). Conversely, the abundances of FLA pathways from *E. siraeum* and *Lachnospira pectinoschiza* were distinctly correlated positively with the HDL-TG level in males (Figure 2A, Data Set 6). We previously demonstrated that flagellins were detected in mouse liver.^11^ Hence, we asked whether flagellins were present in human liver tissues also. Using two different anti-flagellin antibodies raised against flagellins from bacteria of distinct phyla, we detected bands between the size of 26 kDa to 43 kDa in human liver (Figure S3). Other than enterohepatic gateway, the content in gastrointestinal tract can enter through the lymphatic system, that is the drainage of visceral lymph nodes to adipose depots. However, comparing to liver, the flagellin level in retroperitoneal fat was negligible (Figure S3), suggesting the accumulation of flagellins in human liver. Next, two sets of liver protein samples with sizes around 26 kDa and sizes between 26 and 43 kDa were subjected to proteomic analysis using a flagellin proteome database created based on the ones contributing to the FLA pathways in the 500FG cohort.^13^ There were 468 bacterial species detected in the 500FG cohort and 78 of them contributed to the FLA pathways in which 58 species contributed to the readout of GO:0009288 pathway (Figure 2B). The hepatic flagellins at 26 kDa were derived from 23 species in females and 27 species in males, whereas those from 26 to 43 kDa were derived from 21 species in females and 14 species in males (Figure 2C, Data Set 7). These suggest that only a small portion of flagellated bacteria whose flagellins can penetrate through gastrointestinal tract and most of the infiltrated flagellins are in smaller size. Moreover, only 8 of those species were present in at least 50% of individuals in 500FG cohort, including *Eubacterium rectale, E. eligens, E. siraeum, E*. sp. CAG:38, *Lachnospira pectinoschiza, Flavonifractor plautii, Roseburia facecis* and *R. inulinivorans* (Figure 2C, Data Set 7). In summary, *E. rectale* and *R. inulinivorans* were the highly prevalent species in the gut that their flagellins were detected in the liver and positively correlated with HDL-C and ApoA1 levels (Figure 2C, Data Set 7).

**Figure 2.**
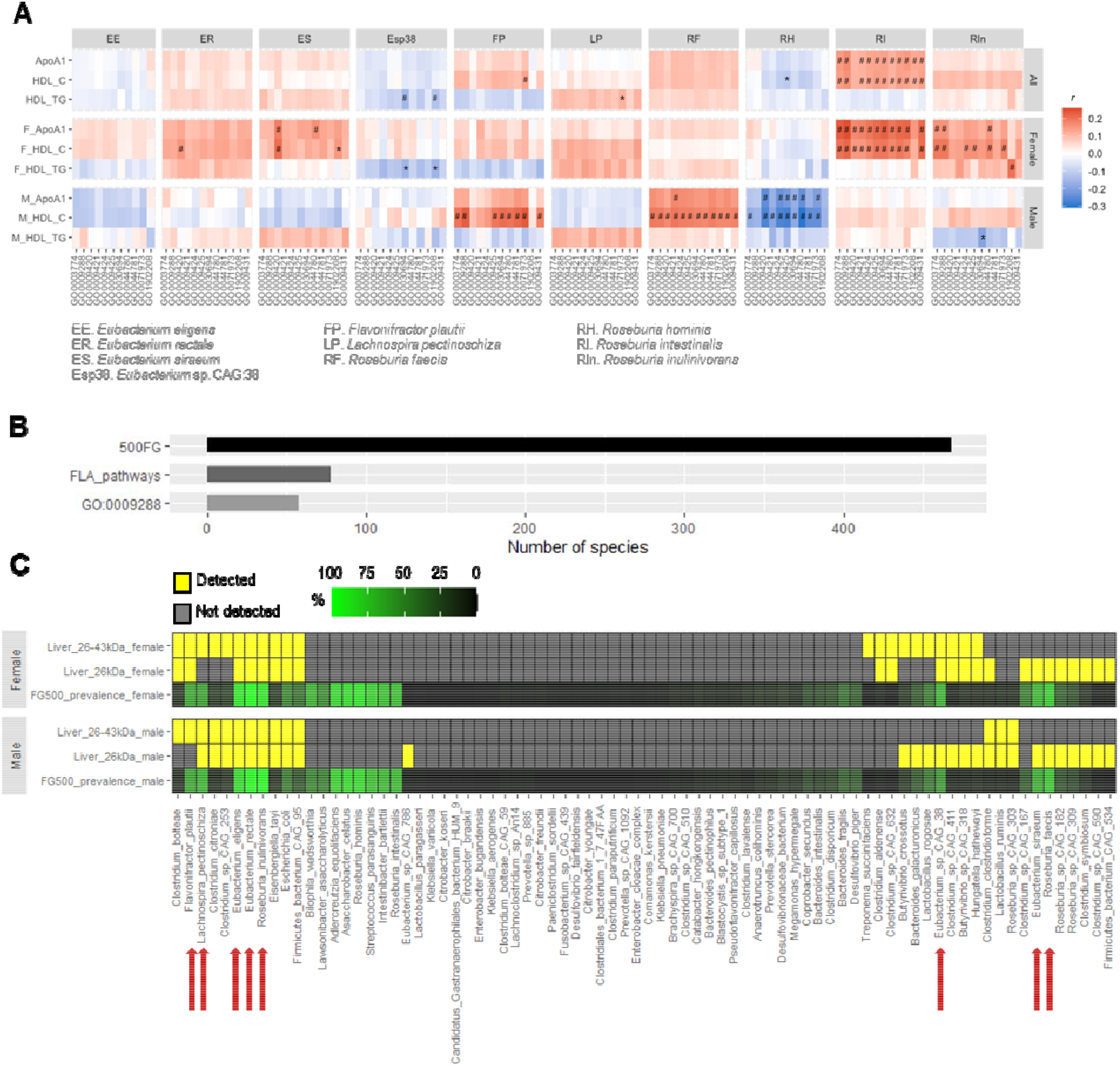
Correlation of flagellated species with HDL-C levels and detection of their flagellins in human livers. (A) Correlations of the abundance of flagellum-related (FLA) pathways from individual flagellated bacteria with ApoA1, HDL-C and HDL-TG levels in males and females of the 500FG cohort. Spearman correlation adjusted with age and BMI for all, and the use of oral contraceptives for females followed by Benjamini-Hochberg procedure for multiple comparison. ^*^*P*<0.05, ^#^*P*<0.05 and *q*<0.2. (B) The numbers of bacterial species detected in the 500FG cohort, contributed to FLA pathways and to the GO:0009288 pathway. (C) The prevalence of bacterial species that contributed to FLA pathway in the 500FG cohort and their presence in human liver tissues (*n*=5 donors per sex). Red arrows indicate the bacteria FLA pathways of which were positively correlated with HDL-C level in (A).

### Different abundance of flagellated bacteria in the cohorts with different ethnicity

We previously observed a positive correlation in overweight males with HDL-C level ≥1 mM but not <1 mM in a small Chinese cohort.^11^ In the 500FG cohort, more than 66% of the males had HDL-C level <1 mM, and, hence, the absence of correlation was consistent with our previous data.^11^ Nonetheless, when we examined another cohort, a Chinese cohort in which 85% of males in the urbanized population had a HDL-C level ≥1 mM (Table 2)^12^, a significant positive correlation between the combined abundance of FLA pathways from *E. rectale* and *R. inulinivorans* and HDL-C level was observed in both males and females (Figure 3A, Data Set 8). In this Chinese cohort, the relative abundances of these 2 species were higher than the 500FG cohort. Moreover, the males in the Chinese cohort showed a higher *R. inulinivorans* abundance (Figure 3B) and stronger correlations between the combined abundance of FLA pathways and HDL-C than females (Figure 3A, Data Set 8). Taken together, it implicates that females in general are more sensitive to these bacteria for increasing HDL-C level. However, when the abundances of these bacteria reach certain threshold, males can also positively respond to them resulting in increased HDL-C level.

**Table 2.** Comparison of females and males from the Chinese cohort.

| Parameter | Female<br>*n* = 193 |  | Male<br>*n* = 154 |  | *P* value |
| --- | --- | --- | --- | --- | --- |
|  | Mean ± S.D. | Range<br>(90<sup>th</sup> percentile) | Mean ± S.D. | Range<br>(90<sup>th</sup> percentile) |  |
| Age | 27.192 ± 9.491 | 18 – 62<br>(43.8) | 26.058 ± 8.164 | 18 – 48<br>(38.0) | 0.002 * |
| BMI (kg/m<sup>2</sup>) | 20.732 ± 2.940 | 14.844 – 31.644<br>(24.21) | 22.058 ± 3.814 | 16.805 – 35.235<br>(27.092) | 0.004 * |
| HDL-C (mM) | 1.43 ± 0.28 | 0.74 – 2.53<br>(1.80) | 1.26 ± 0.31 | 0.54 – 2.52<br>(1.58) | 6.37 × 10<sup>-6</sup> * |
| LDL-C (mM) | 2.54 ± 0.71 | 1.01 – 5.45<br>(3.40) | 2.67 ± 0.77 | 0.95 – 5.04<br>(3.62) | 0.191 |
| TC (mM) | 4.02 ± 0.78 | 2.55 – 7.18<br>(4.92) | 4.00 ± 0.85 | 2.28 – 6.65<br>(5.10) | 0.810 |
| TG (mM) | 0.99 ± 0.68 | 0.35 – 7.62<br>(1.57) | 1.27 ± 0.99 | 0.39 – 5.74<br>(2.19) | 0.015 * |

\* *P* < 0.05, two-sided *t* test

**Figure 3.**
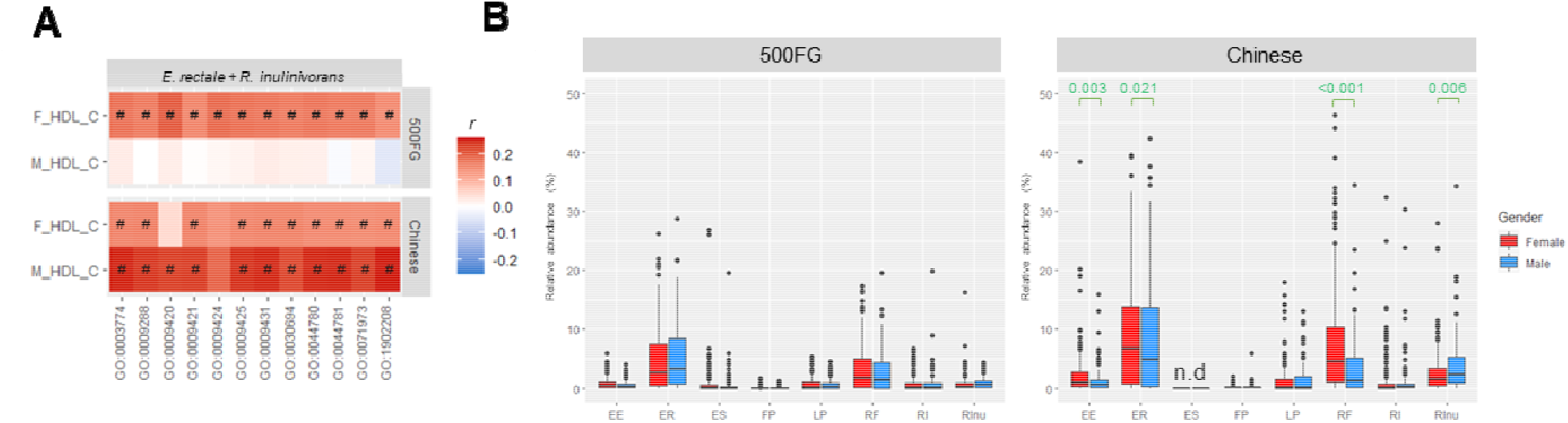
Comparison of correlations between the 500FG and Chinese cohorts. (A) Correlation of HDL-C level with the summed abundances of FLA pathways from *Eubacterium rectale* and *Roseburia inulinivorans*, and (B) the relative abundances of different flagellated bacteria in males and females from the 500FG cohort and the Chinese cohort. Spearman correlation adjusted with age and BMI for all, and the use of oral contraceptives for females in 500FG cohort followed by Benjamini-Hochberg procedure for multiple comparison in (A). ^*^*P*<0.05, ^#^*P*<0.05 and *q*<0.2. n.d., not detected. The full names of the bacteria are listed in Figure 2.

### Detections of TLR5 but not NLRC4/NAIP in human liver

Hepatic TLR5 was responsive to gut bacteria-derived flagellin in mice, and treatment with flagellin stimulated ApoA1 production in human hepatocytes.^11^ We then explored whether TLR5 protein in liver played a role in the sex difference in the sensitivity to flagellins in terms of the favorable cardiometabolic outcome. TLR2, TLR5, TLR9 and TLR10 protein were detectable in both female and male liver samples while TLR6, TLR7 and TLR8 were undetectable (Figure 4A). We also examined the aortic samples as comparison, and TLR5 were undetectable in both sexes (Figure 4A). Because we used the whole liver for proteomic analysis, hepatic immune cells could contribute to the proteomic readout. Nonetheless, instead of the full spectrum of TLR family, only certain TLRs were detected, suggesting that the proteomic readouts reflected the protein abundance in the hepatocytes rather than the less populated non-parenchymal cells. In terms of TLR5, it was detected in 4 out of 5 and 3 out of 5 liver samples in females and males, respectively (Figure 4B), and its abundance was higher in males than females, suggesting TLR5 in females might be more sensitive to the infiltrated flagellins (Figure 4B). Other than TLR5, flagellin can engage with NAIP/NLRC4 to activate inflammasome, but neither NAIP nor NLRC4 was detectable in these 10 liver samples (Figure S4). At transcription level, *NLRC4* mRNA was significantly higher in male liver samples (Figure 4C). Moreover, *TLR5* showed strong negative correlation with *NLRC4* and positive correlation with *NAIP* in female livers (Figure 4C). Conversely, in male livers, both strong correlations were absent (Figure 4C). The sex difference in the relationship between *NLRC4* and *TLR5* transcription requires further study.

**Figure 4.**
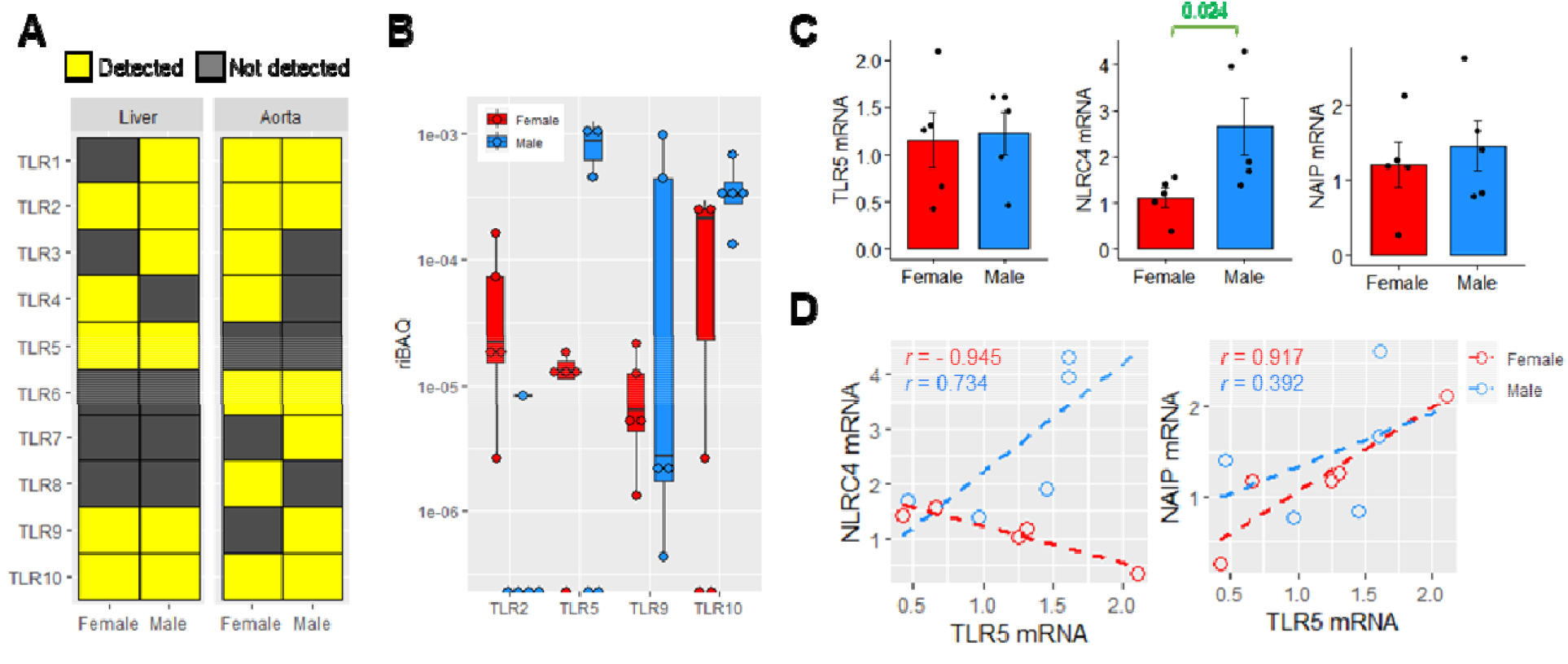
Sex difference in the expression of flagellin-engaging receptors. (A) The detection of various TLRs in human livers and abdominal aortic lesions in females and males (*n*=5 per sex for livers, *n*=3 per sex for aortic lesions). (B) The relative intensity-based absolute quantification (riBAQ) of different TLRs in human liver samples (*n*=5 per sex). (C) Relative mRNA expression of *TLR5, NAIP* and *NLRC* in female and male livers (*n*=5 females, *n*=6 males). Two-sided unpaired *t* test. (D) Correlation of the mRNA expressions of *NAIP* and *NLRC4* with *TLR5* in livers. Pearson correlation.

## Discussion

The detection of bacterial proteins in our body has been reported in sepsis or infectious diseases only but rarely in non-infectious conditions. Here, we show that bacterial flagellins were detected in human liver and the abundances of FLA pathways from *Eubacterium rectale* and *Roseburia inulinivorans* in feces were highly associated with HDL-C level. The liver samples in this study were derived from patients without active infections, and the detection of flagellins in liver implicates a tolerance for the penetration of bacterial proteins in the body. Commensal bacteria are known to evolve various mechanisms to evade host surveillance. Certain flagellated bacteria or flagellins escape the host’s mucosal immunity and the host eventually evolves mechanisms benefitting from such evasion also. Indeed, several species of gut bacteria, including *E. rectale*, have been found to be co-diversified with human, which means a shared evolutionary history resulting in a reinforcement on commensal relationship across generations.^17^ The flagellins detected in liver resulting in ApoA1 production might represent an outcome of the co-diversification. A recent study reported that a new subclass of flagellins, termed silent flagellins, which are largely produced by members from Lachnospiraceae family, have a lower affinity to dimeric TLR5 compared with well-studied, potent flagellins such as those from the pathogenic *Salmonella enterica* yet retain their efficacy to stimulate TLR5 at high concentrations.^18^ *E. rectale* and *R. inulinivorans* belong to this family. It is likely that such silent property allows the flagellins from commensal bacteria to bypass the intestinal mucosal checkpoint but triggers hepatic TLR5 once they accumulate in the liver.

A limitation of this study is the different sources of human samples, such that we were unable to compare the detections of flagellated bacteria in the gut and hepatic flagellins in the same individuals. The lack of fecal samples for the examination of actual flagellin protein abundances prevented us to validate whether the undetected flagellins from certain species were due to the absence of the bacteria or the impermeability to the host. The age difference in the samples is another limitation, the average ages of the 500FG cohort and tissue samples are 27.4 and 68.4, respectively. Aging and menopause may affect the mucosal immunity altering the penetration of flagellins. Nonetheless, despite the age difference, the proteomic experiments also identified the types of bacteria showing positive correlations between fecal FLA pathways and circulating ApoA1 level.

Through a multi-omic approach to investigate the sex difference in the cardiometabolic role of gut microbiota, we identified that commensal flagellated bacteria abundant in the gut could explain the host HDL-C and ApoA1 levels, which, along with the detection of their flagellins in liver tissues, confirms our previous in vitro observations on human primary hepatocytes that flagellins can stimulate Apoa1 production by engaging TLR5. We believe that this interaction between gut microbiota and host will inspire us a new design in treatment for increasing ApoA1 and HDL-C levels. The development of orally stable TLR5 agonists to stimulate ApoA1 production in the liver is worth pursuing.

## Acknowledgements

None

## Supplemental Material

Supplemental Methods

Tables S1-3

Figures S1-4

Data Sets 1-8

Major Resources Table

